# Type IV Secretion System Drives Lipid Mixing

**DOI:** 10.64898/2026.01.14.699567

**Authors:** David Chetrit, Craig R. Roy, Erdem Karatekin

## Abstract

Type IV secretion systems (T4SSs) are versatile molecular machines used by bacteria to secrete protein effectors into host cells, promoting pathogenesis, and to transfer DNA between bacteria through conjugation, driving horizontal gene transfer. Most, like Dot/Icm of the pathogen *Legionella pneumophila* (*L. pneumophila)* or *Escherichia coli* (*E. coli)* RK2, are primed for substrate delivery only upon contact with a target membrane, but mechanisms are unknown. A pilus could bind a receptor to initiate priming, but many T4SSs, especially those that deliver effectors, lack a pilus. Here, we present evidence that T4SSs are primed by direct contact with target membrane lipids. Combining fluorescence assays with genetics and biochemistry, we found that Dot/Icm drives lipid exchange between bacterial cells and between bacteria and synthetic membranes containing only lipids. Lipid exchange requires membrane contact but does not require ATP hydrolysis or even full complex assembly. Minimally, the outer membrane core complex protein DotG needs to be present in at least one of the apposed membranes. We similarly observed lipid mixing with the simpler *E. coli* RK2 T4SS, where we could follow lipid mixing and plasmid transfer simultaneously. We found that lipid mixing always preceded or accompanied plasmid transfer, suggesting it may be part of the contact-dependent priming mechanism. Lipid mixing was inhibited or promoted by lipids that inhibit or promote membrane fusion, respectively. Lipids inhibiting lipid mixing also inhibited substrate transfer. Together, our results suggest that initial contact between DotG outer segments and target membrane lipids promotes lipid mixing as part of the mechanism that primes T4SS for substrate translocation.

**Highlights:**

1. The *Legionella pneumophila* Dot/Icm secretion system requires contact with a target membrane for effector translocation, but how membrane contact primes the machinery for this is unknown.
2. We found that Dot/Icm drives contact-dependent lipid mixing between bacteria or between bacteria and liposomes, showing that Dot/Icm priming does not require a protein receptor.
3. Lipid mixing also occurs with the simpler Type IVA system (*Escherichia coli* RK2) during conjugation, suggesting lipid mixing is a general feature of Type IV Secretion System function.
4. The core complex protein DotG is sufficient to drive lipid mixing, suggesting DotG-target membrane interactions may destabilize the target membrane.

## INTRODUCTION

The bacterium *L. pneumophila* delivers protein effectors into host cells through a secretion system called Dot/Icm to avoid lysosomal degradation and create a niche for replication. Although *L. pneumophila*’s natural hosts are freshwater amoebae, it is also an opportunistic human pathogen that causes a severe pneumonia known as Legionnaires’ disease^1,2^. Dot/Icm is essential for infection and pathogenesis: it engages with the plasma and vacuolar membranes of host cells to deliver over 300 different effectors into the host cytosol^3–5^ (Fig. 1A). These effectors hijack host mechanisms to support bacterial survival and replication within specialized vacuoles, which expand through the acquisition of endoplasmic reticulum (ER)-derived membranes^6,7^. Interestingly, the same Dot/Icm machinery can mediate horizontal gene transfer among bacterial cells through conjugation, promoting the spread of fitness-enhancing traits such as antibiotic resistance and virulence factors (Fig. 1A-inset)^8^. Remarkably, the Dot/Icm system has also been shown to translocate effectors directly between bacterial cells in a conjugation-like manner, confirming its dual role in both inter-kingdom pathogenesis and intra-bacterial communication^9^. Dot/Icm lacks a stable pilus, and no specific receptor has been identified^3,10–12^. Despite their importance, how Dot/Icm and other Type IV secretion systems (T4SSs) are primed for substrate delivery is not known^13–18^.

**Figure 1.**
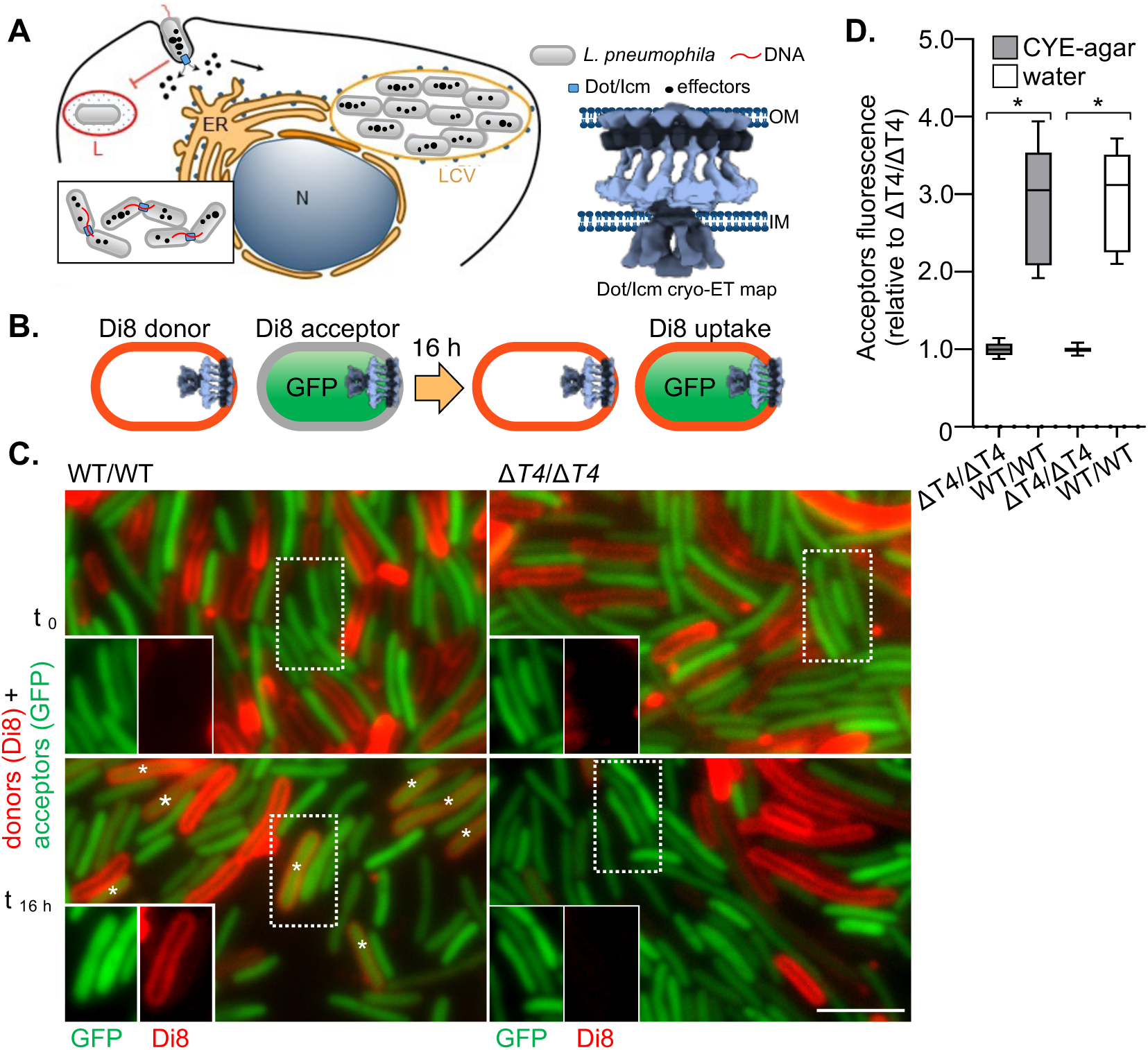
Dot/Icm system promotes lipid mixing. **A.** Left. Upon interacting with the host cell membrane, *L. pneumophila* delivers effectors through the Dot/Icm T4SS into the host cytoplasm. These effectors prevent the degradation of internalized bacteria in lysosomes (L) and instead facilitate bacterial replication within a specialized Legionella-containing vacuole (LCV). N: nucleus, ER: endoplasmic reticulum. Inset (bottom left). The Dot/Icm preserves the ancestral capacity of T4SSs to deliver DNA to other bacteria through a conjugation-like mechanism. Right-Cryo-electron tomography map of the Dot/Icm secretion apparatus embedded within the bacterial envelope, spanning the inner (IM) and outer membranes (OM)^5,48,69^. **B-C.** *L. pneumophila* donor cells, either wild type (WT) or an isogenic mutant lacking all *dot/icm* genes (Δ*T4*), were labeled with the membrane dye Di8, grown for 16 h on solid media (CYE-agar), and co-incubated with acceptor cells expressing cytosolic GFP. Imaging revealed that GFP-positive acceptor cells took up the donor membrane dye (*), indicating lipid mixing between donors and acceptors. A dashed square indicates an inset of the acceptor cells. Scale bar: 3 µm **D.** Lipid mixing between *L. pneumophila* cells is Dot/Icm-dependent and does not require bacterial growth. Wild-type (WT) and Δ*T4SS* (Δ*T4*) donor and acceptor *L. pneumophila* strains were labeled with the membrane dye Di8 and either co-incubated on solid media or pelleted and incubated in water for 16 h. For the solid media condition, data represent the means of five independent replicates (n = 5056 WT and 4248 ΔT4 cells; two-sided Student’s t-test, *p < 0.0008*). For the water incubation condition, data represent three independent replicates (n = 1012 WT and 950 Δ*T4* cells; two-sided Student’s t-test, *p < 0.0001*).

T4SSs form a large family, broadly divided into two subfamilies, based on genetic and structural similarities. Type IVA secretion systems (T4ASSs) are simpler, containing 8-12 protein components^18,19^. Well-characterized examples include the *Agrobacterium tumefaciens* (*A. tumefaciens*) VirB/VirD4 system, the *E. coli* R388, pKM101 conjugation systems, and the RK2 plasmid, which encodes a closely related T4ASS^20^. RK2 is a broad-host-range IncP plasmid capable of replicating in various Gram-negative bacteria, making it a valuable tool in genetic engineering^21^. The Dot/Icm system is part of the more complex Type IVB systems (T4BSSs), which contain approximately twice as many components. It is chromosomally encoded by pathogens such as *L. pneumophila*, which is the best-studied example of a T4BSS^22^. Several components are conserved between T4ASSs and T4BSSs, suggesting some shared mechanisms. Notably, the core complex proteins VirB10 and DotG show clear homology, as do the three ATPases: the VirD4/DotL substrate receptors (also called Type IV coupling proteins or T4CPs), VirB4/DotO, and VirB11/DotB. Several structural models of T4SSs in their inactive (non-translocating) states have identified component locations. Cryo-electron tomography (Cryo-ET) analysis revealed that the *L. pneumophila* Dot/Icm system forms a complex consisting of an outer membrane core complex (OMCC) and an inner membrane complex^5^ (IMC) (Fig. 1A, right). The OMCC includes two distinct curved protein layers, one just beneath the outer membrane (OM) and one in the periplasm, with additional unresolved electron density extending above the OM. These layers are composed of at least five subunits: DotC, DotD, DotF, DotG, and DotH. The cytoplasmic complex includes ATPases DotO and DotB, homologs of which are shared with T4ASSs and are required to energize substrate delivery^3,23–32^.

Dot/Icm-mediated effector translocation requires contact with a target membrane, as constitutive secretion is not observed. The proteins DotA and IcmX assemble a periplasmic protochannel in the center of the secretion apparatus. Upon host membrane contact this protochannel undergoes a conformational rearrangement that results in an opening of the channel and extension of the apparatus through the outer membrane. How the Dot/Icm system senses host membranes to trigger this conformational change that activates the secretion system remains unknown. Here we define priming as contact-triggered conformational changes that prepare the T4SS for substrate delivery. After priming, the T4SS is in a state that can deliver substrate, provided further activation steps are satisfied, including those that are mediated by T4 coupling proteins and ATPases^33^. While binding to a specific receptor protein could serve as a priming trigger, the *L. pneumophila* Dot/Icm mediates substrate translocation across a broad range of target membranes, from bacteria to amoebae and human cells, and no such receptor has been identified for the Dot/Icm system, indicating that a specific proteinaceous receptor may not be necessary^34^. Another possibility for Dot/Icm priming involves a structural projection that senses nearby membranes. Yet, unlike Type III or VI secretion systems, Dot/Icm and most other T4BSSs lack a stable needle-like projection^3,10,27,32^. Consistently, recent work found that *L. pneumophila* forms Dot/Icm-dependent tight contact sites with target membranes in infected *Acanthamoeba castellanii, Dictyostelium discoideum,* and HeLa cells^12^. These sites often exhibit hourglass-shaped deformations at the *Legionella*-containing vacuole (LCV) and bacterial OM, suggestive of lipidic continuity. Given that a proteinaceous receptor is unlikely to be involved, here we hypothesized that direct interactions of Dot/Icm with target membrane lipids may lead to its priming for substrate transfer. We further reasoned that if this is the case, then lipid exchange might occur between the tightly apposed membranes. Supporting this idea, we found Dot/Icm-and contact-dependent lipid mixing between *L. pneumophila* cells. Lipid mixing was also observed during RK2-dependent conjugation in *E. coli*, suggesting it may be a conserved feature across at least some T4ASSs and T4BSSs. In *E. coli*, where we could monitor lipid mixing and substrate transfer simultaneously, lipid mixing always preceded or coincided with substrate delivery but never followed it. Contact with artificial lipid membranes were sufficient to trigger Dot/Icm-dependent lipid mixing, suggesting that membrane lipids alone can prime the machinery, which would explain the host promiscuity of Dot/Icm systems.

## RESULTS

### The Dot/Icm system drives lipid mixing between bacteria

Based on prior findings showing that the Dot/Icm system can actively support conjugation and even translocation of effectors directly between bacteria^8,9^, we tested whether Dot/Icm also mediates lipid exchange between bacterial cells. In these experiments, the membranes of donor *L. pneumophila* were labeled with the lipophilic marker Di-8-ANEPPQ (Di8), which, like other styryl dyes, is essentially non-fluorescent in aqueous solution but becomes highly fluorescent once inserted into a membrane^35–37^ (Fig. 1B). After labeling, donor cells were washed to remove excess dye. Acceptor *L. pneumophila* had unlabeled membranes but expressed cytoplasmic GFP so they could be differentiated from donors. Wild-type (WT) or Δ*T4SS* (Δ*T4*) donor and acceptor cells were mixed and placed on solid media (CYE-agar) and visualized 16 h later using fluorescence microscopy (Fig. 1C). We found that Dot/Icm promoted lipid mixing, as the level of fluorescence in acceptor membranes was higher in wild-type donor/wild-type acceptor (WT/WT) pairs compared to Δ*T4*/Δ*T4* controls (Fig. 1C). In the Δ*T4*/Δ*T4* condition, few acceptor cells acquired detectable labeling, likely due to some partitioning of the dye between membranes and the medium. To quantify lipid transfer across the entire acceptor population, fluorescence was measured from randomly chosen acceptor cells across multiple fields of view, ensuring unbiased sampling; on average, WT/WT showed ∼3-fold higher fluorescence than Δ*T4*/Δ*T4* (Fig. 1D).

Bacterial growth could possibly affect dye uptake. To test if this is the case, donor and acceptor cells were pelleted and incubated in water for 16 h, a condition that prevents growth but permits cell-to-cell contact and survival (Fig. 1D)^38–40^. Even under these conditions, Dot/Icm promoted lipid mixing, suggesting that lipid exchange occurs independently of bacterial proliferation.

### Lipid mixing requires the Dot/Icm system and cell-to-cell contact

To better characterize the temporal dynamics of lipid mixing, we performed time-lapse imaging over 16 h using Di8-labeled *L. pneumophila* donor cells incubated with either WT or Δ*T4* acceptor cells on an agarose pad (Fig. 2A,B). Under these conditions, only WT acceptor cells that were in direct contact with WT donor cells acquired Di8 fluorescence, with the fluorescence signal in the acceptor cell spreading from the contact site. Such sites were predominantly at the bacterial poles, where Dot/Icm complexes localize^3,10,12^, and reaching a plateaus after ∼8 h (Fig. 2C). In contrast, Δ*T4* acceptors incubated with Di8-labeled Δ*T4* donors showed no detectable fluorescence, indicating that lipid exchange requires a functional Dot/Icm system. No fluorescence increase was observed in WT recipients that were not in contact with WT donors or in Δ*T4* donor-Δ*T4* acceptor pairs, underscoring the requirement for both Dot/Icm activity and cell-cell contact.

**Figure 2.**
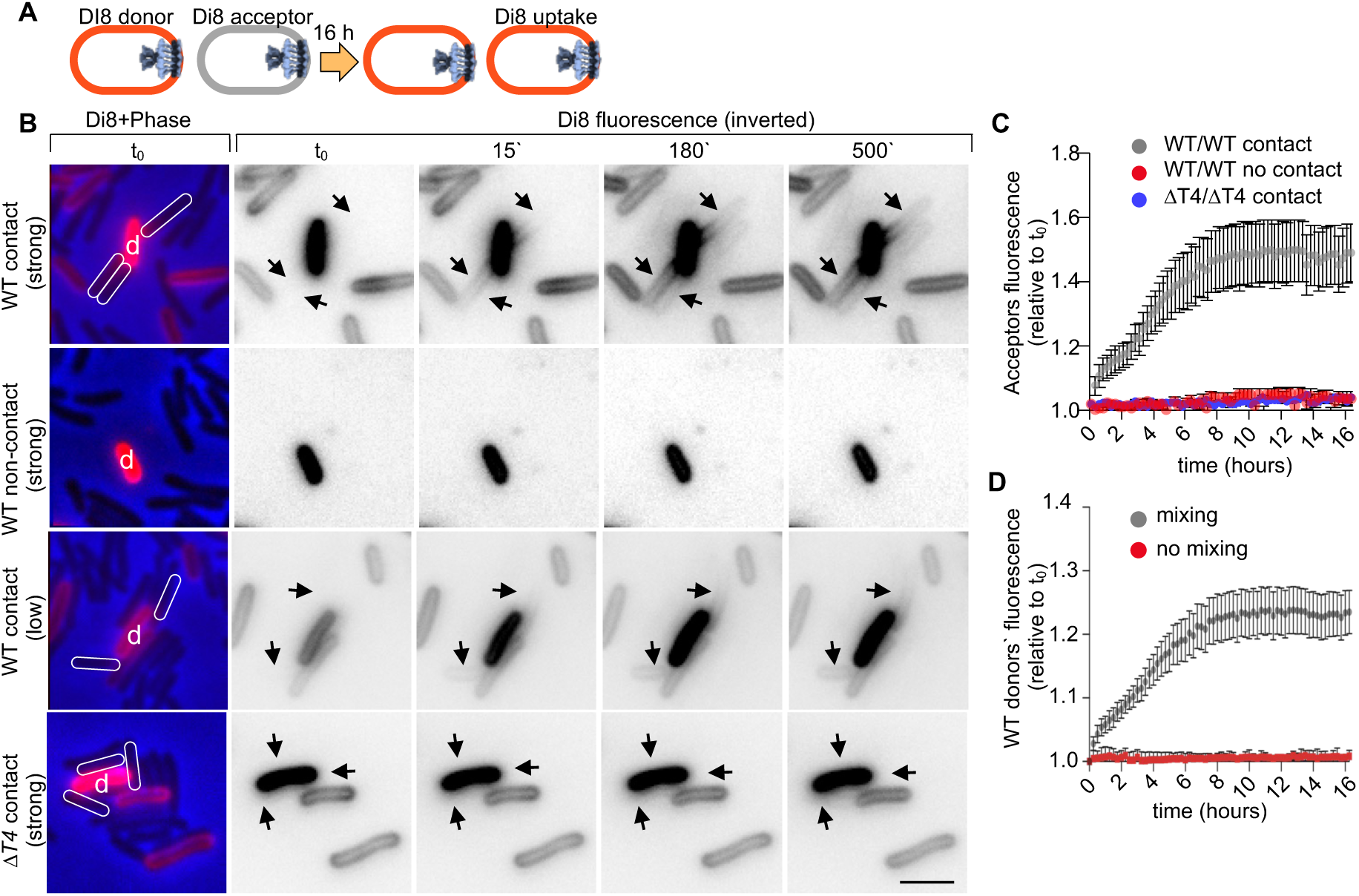
Time-lapse imaging shows lipid mixing depends on Dot/Icm and cell-cell contact. **A**. Wild-type (WT) *L. pneumophila* donors labeled with Di8 (red) were mixed with unlabeled WT or Δ*T4* acceptors and imaged under an agarose pad. **B.** Phase contrast imaging of the first frame was used to identify acceptors. At subsequent time points, Di8 fluorescence intensity was monitored. Strongly (strong) or weakly (low) labeled donor cells are marked with “d,” and contacting acceptors cells are indicated by arrows and outlines. Only WT acceptors in direct contact with donor cells showed Di8 uptake, with the fluorescence signal spreading from the site of contact. Δ*T4* acceptors did not acquire any Di8 fluorescence. Scale bar: 2 µm. **C.** Average change in Di8 fluorescence over time for contacting acceptors. Lipid exchange occurred between WT donors and WT acceptors (WT/WT) but not between non-contacting WT donors or between *ΔT4* donors and Δ*T4* acceptors (Δ*T4*/Δ*T4*). (n=30 cells per condition, 3 independent replicates). Error bars represent the standard error of the mean (SEM). **D.** Average change in Di8 fluorescence for WT donors over time. Lipid mixing was accompanied by dequenching of Di8 and increased fluorescence intensity in most donor cells. No significant change was observed when lipid exchange did not occur. (n=25 donor cells per condition, 3 independent replicates). Error bars represent SEM.

Lipid transfer was frequently accompanied by dequenching of the Di8 dye in donor membranes, resulting in increased fluorescence intensity in donor cells (e.g. Fig. 2B, 3^rd^ row). This phenomenon is consistent with the known properties of Di8, and indicates that in some donor cells its concentration reached self-quenching levels prior to interacting with acceptor cells^41^. Upon lipid mixing with an acceptor cell, Di8 transfer to the acceptor membrane dilutes it in the donor membrane, leading to donor fluorescence dequenching. When present, dequenching thus serves as an additional indication of lipid exchange between donor and acceptor membranes. Time-lapse microscopy of donor cells showed a progressive increase in Di8 fluorescence that plateaued after approximately 8 h (Fig. 2D), consistent with kinetics of transfer observed above. Together, these results demonstrate that lipid exchange requires both a functional Dot/Icm secretion system and direct physical contact between donor and acceptor cells.

### Lipid mixing is modulated by membrane fusion regulators

The simplest mechanism that could cause lipid mixing is direct membrane continuity between the donor and acceptor membranes, i.e. hemifusion (when only the proximal leaflets have undergone fusion, but not the distal ones) or fusion (when both leaflets of the donor and acceptor membranes are continuous). Both are inhibited by positive curvature lipids in the proximal leaflets and promoted by addition of negative curvature lipids^42^. To examine whether Dot/Icm-mediated lipid mixing involves membrane fusion intermediates, we tested the effects of oleic acid (OA) or lysophosphatidylcholine (LPC), lipids known to modulate membrane curvature^43^ (Fig. 3A). OA, a negative curvature lipid, stabilizes the membrane stalk intermediate and facilitates (hemi)fusion, whereas LPC, a positive curvature lipid, destabilizes this intermediate and inhibits (hemi)fusion when incorporated into the outer leaflet (Fig. 3A). To assess the impact of these membrane curvature modulators on lipid mixing, Di8-labeled *L. pneumophila* donor cells were co-cultured with GFP-expressing acceptor cells for 16 h on solid media in the presence of either OA or LPC, and then imaged (Fig. 3B). Imaging showed that OA enhanced lipid transfer to acceptor cells, while LPC markedly reduced it (Fig. 3C). OA treatment increased lipid mixing by ∼75% relative to untreated wild-type cells, whereas LPC reduced lipid transfer by ∼25% (Fig. 3D). These treatments had no effect on the small amount of lipid exchange observed between Δ*T4* donor-Δ*T4* acceptor pairs (Fig. 3D). Overall, these results indicate that positive- or negative-curvature lipids that are known to modulate membrane fusion significantly affect Dot/Icm-mediated lipid mixing, suggesting the involvement of membrane fusion intermediates.

**Figure 3.**
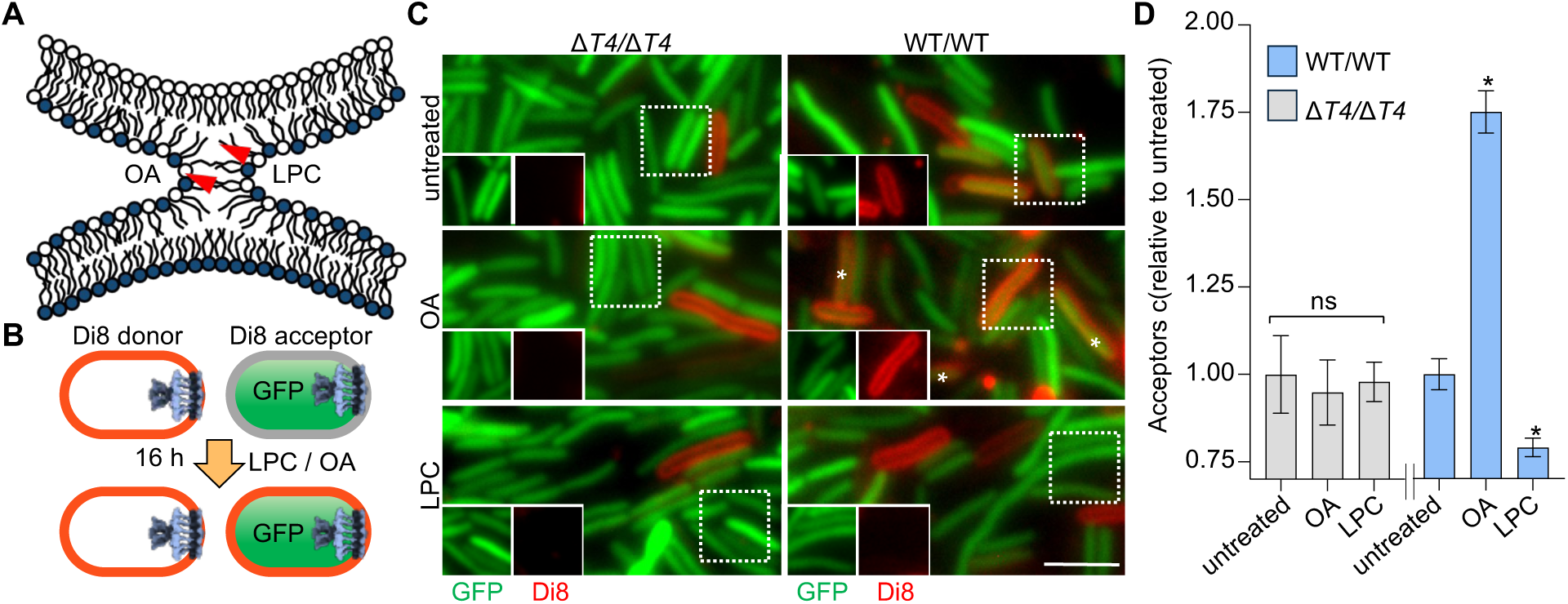
Lipid mixing is affected by membrane fusion modulators. **A**. A membrane stalk is thought to constitute a universal intermediate that later develops into a hemifused and then a fused state. Because the stalk has net negative curvature, it is stabilized by negative curvature lipids such as oleic acid (OA) and conversely destabilized by positive curvature lipids such as lysophosphatidylcholine (LPC). Thus, OA facilitates whereas LPC inhibits fusion when added to the outer leaflets. **B**. *L. pneumophila* donors labeled with Di8 were grown with acceptors expressing GFP (green) for 16 h with OA or LPC on solid media and imaged. **C-D**. Representative micrographs and the amount of donor membrane dye transferred to acceptor cells, expressed as the mean pixel value of acceptors normalized by the intensity of untreated cells. For Δ*T*4/Δ*T*4, n=616, 978, and 1101 cells were analyzed for untreated, OA, and LPC conditions, respectively. For *WT/WT*, n=1400 cells were analyzed for each condition. Two biological replicates. * *p<0.05*, two tailed student’s t-test. Scale bar, 3 µm.

### Lipid mixing precedes or occurs concomitantly with substrate translocation

To define the temporal relationship between lipid mixing and substrate delivery, and to determine whether lipid mixing is conserved across different types of T4SSs, we investigated the conjugative transfer of the RK2 plasmid, encoding a canonical *E. coli* T4A conjugation system. The plasmid was engineered to express GFP (RK2-GFP) to allow detection of successful conjugation events into acceptor cells. Donor cells were labeled with NileRed, a lipophilic dye that fluoresces strongly in lipid-rich, but not in the aqueous, environment, while acceptor cells were stained with Hoechst to distinguish them from donors. This assay can distinguish four possible types of acceptors: those that received neither lipid nor plasmid (N); those that received only lipid (L); those that received both lipid and plasmid (LP); and those that received only plasmid but no lipid (P). After 16 h of co-incubation on solid media (LB-agar), no lipid or plasmid transfer was observed when donors carried the non-conjugative control plasmid pJB1806-GFP (Fig. 4A-B), confirming that a functional T4SS is required for lipid mixing. By contrast, robust lipid mixing and plasmid transfer was evident when donors carried the RK2-GFP plasmid. The NileRed lipid labeling was dim in pJB1806-GFP donor cells, but not in RK2-GFP ones, likely due to a lack of dequenching that would result from lipid mixing with acceptor membranes in the former (cf. Fig. 2). For cells expressing RK2, we quantified the statistics of the four acceptor types in Fig. 4C. When donors expressed RK2-GFP, 84% of acceptor cells exhibited NileRed fluorescence, but no GFP, i.e. they received only lipids (L). Fourteen percent of acceptor cells expressed GFP and had NileRed labeled membranes indicating successful RK2-GFP plasmid conjugation and lipid transfer (LP). Only 2% of the acceptors remained unlabeled (N). Remarkably, all GFP-positive acceptors were also NileRed-positive, but there were no GFP-positive acceptors that were negative NileRed (P). These results indicate that T4SS-dependent lipid mixing always accompanied substrate translocation, occurring before or during substrate transfer. If substrate transfer occurred in the absence of lipid mixing, we would have observed some GFP-positive acceptors without any lipid mixing, which was not the case.

**Figure 4.**
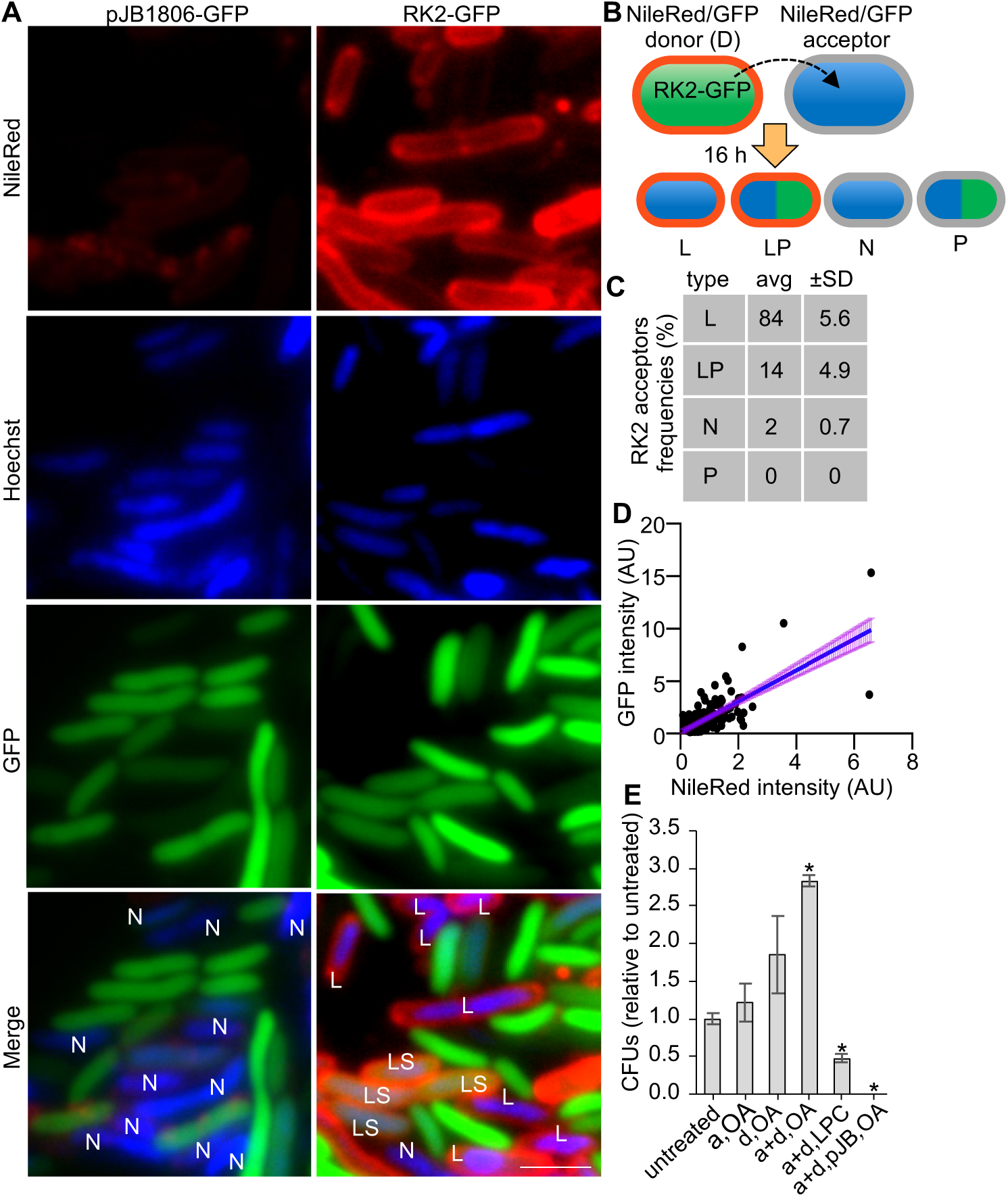
Lipid mixing in RK2-conjugating *E. coli* precedes or occurs concomitantly with substrate transfer. **A-B**. *E. coli* donor cells carrying either the control plasmid pJB1806-GFP or the RK2-GFP plasmid (green) were labeled with NileRed and incubated for 16 hours with Hoechst-labeled *E. coli* acceptor cells (blue). Merged image show different classes of acceptor cells as labeled: L: lipid transfer alone; P: plasmid transfer alone; LP: lipid+plasmid transfer; N:no transfer. Scale bar: 4 µm. **C.** Statistics of lipid and plasmid acquisition by acceptor cells from RK2-GFP expressing, NileRed labeled donors. 1089 cells from 2 biological replicates. **D.** Correlation between lipid exchange and plasmid transfer. Total NileRed intensity and total GFP intensity were measured in individual acceptor cells. Linear regression (blue line, R^2^=0.52) is shown together with 95% confidence bands (purple). A total of 212 acceptor cells were included**. E**. Modulation of membrane fusion impacts *E. coli* conjugation. Donors (d), acceptors (a), or both (d+a) were incubated with OA (facilitates membrane fusion) or LPC (inhibits membrane fusion, see Fig. 3). pJB is the control plasmid pJB1806. n=3 replicates. * p<0.05 two-tailed t-test.

We reasoned that if lipid mixing is required before or during substrate transfer, then the two transfer rates may be correlated. To test for this, we plotted the NileRed and GFP intensity of individual acceptors, as shown in Fig. 4D. We found that an acceptor that acquired high (low) amount of lipid labels was also likely to express high (low) levels of GFP. This observation suggests that lipid and substrate transfer rates are correlated.

Is it possible that substrate transfer is underestimated because of a delay in GFP expression in acceptor cells following plasmid acquisition? This is unlikely to explain the results, because analysis was performed 16 hours after co-incubation, sufficient for GFP expression and accumulation^44,45^.

Our results above are best explained by lipid mixing being a prerequisite for substrate transfer. If this is correct, then introduction of lipids promoting or inhibiting lipid mixing should have the same effect on substrate transfer (Fig. 3). To test this idea in a label-independent manner, we incubated cells either with fusion promoting OA or fusion-inhibiting LPC and measured the conjugation efficiency of RK2. OA increased (⁓3 fold) and LPC decreased (⁓2 fold) colony-forming unit (CFU) counts of transconjugants (Fig. 4E), supporting the idea that lipid mixing is a prerequisite for substrate transfer.

Up to here, our results support the idea that lipid mixing is part of the priming process that is required to initiate or facilitate plasmid transfer. Some lipid mixing occurs in the absence of substrate transfer, possibly through transient contacts that do not lead to successful substrate translocation.

### Lipid mixing requires DotG, but no T4SS ATPases

To identify which components of the Dot/Icm T4SS are required for lipid mixing, we performed lipid mixing assays using *L. pneumophila* donor and acceptor strains expressing either wild-type or mutant Dot/Icm systems (Fig. 5A, B). The mutants included: DotB_E191K_, an ATPase-deficient mutant locked in the ATP-bound conformation that forms a fully assembled but inactive system; Δ*dotA*, which assembles the OMCC but does not have the central DotA/IcmX protochannel and fails to recruit cytosolic components; and Δ*dotG*, which cannot assemble the OM dome component DotG but retains the remainder of the OMCC^3,10,26,27,32^. Quantification of Di8 fluorescence in acceptor cells (Fig. 5C), normalized to ΔT4/ΔT4 controls, showed reduced but significant lipid mixing in combinations where only one strain expressed the full Dot/Icm system or just the OMCC, including WT with Δ*dotA* or Δ*T4*, Δ*dotA* with WT or Δ*dotA,* and Δ*T4* with WT. Thus, full assembly of the apparatus is not required for lipid mixing; the presence of the OMCC in a single interacting membrane is sufficient. Consistent with this, the ATPase-deficient DotB_E191K_ mutant supported lipid mixing at wild-type levels, indicating that ATP hydrolysis is also dispensable. By contrast, the Δ*dotG* mutant was completely defective, and complementation with plasmid-expressed DotG restored lipid transfer. Remarkably, expression of DotG alone in a T4SS-null background was sufficient to drive lipid mixing when present in either donors or acceptors, demonstrating that DotG is both necessary and sufficient to mediate intercellular lipid exchange.

**Figure 5.**
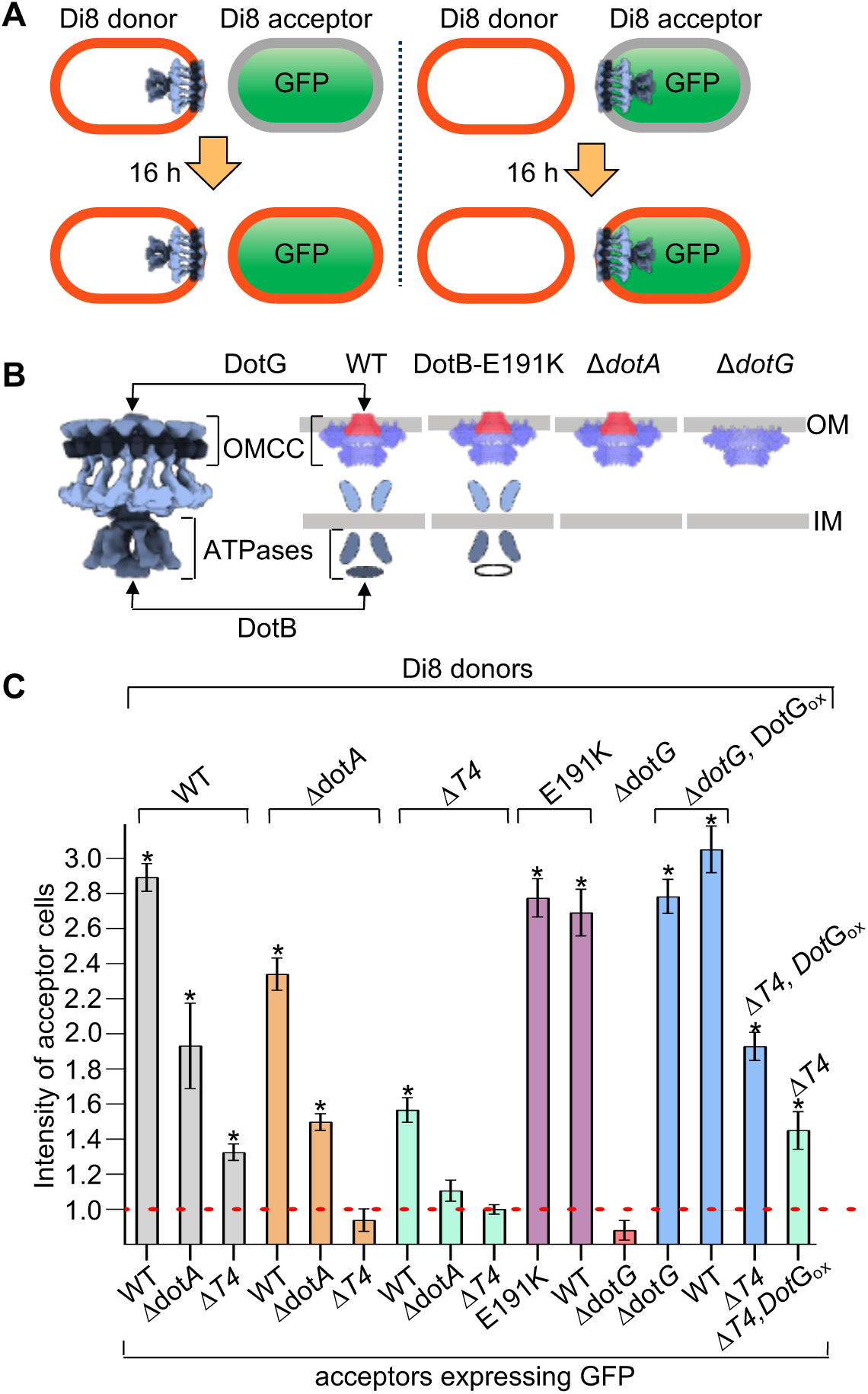
DotG is required for lipid mixing while T4SS ATPases are dispensable. **A.** *L. pneumophila* donors labeled with Di8 were grown with acceptors expressing GFP (green) for 16 h. Strains expressing wild type or mutant forms of the Dot/Icm systems (see B) were used as donor or acceptor cells. **B**. Structures of Dot/Icm mutants used are shown schematically. ΔdotA (ΔA) assembles solely the OMCC, DotB_E191K_ produces a fully assembled apparatus that is ATP-locked and unable to translocate effectors, ΔdotG (ΔG) assembles solely the OMCC without DotG (red). **C**. Lipid mixing in *L. pneumophila* determined as the total acceptor Di8 intensity relative to ΔT4/ΔT4 (red dashed line). Lipid mixing was performed on solid media and measured 16 h later. ΔT4,Gox expresses DotG from a plasmid in a Δ*T4* background. 3 replicates, 3000 cells per condition. *\*p < 10^-5* (two-tailed Student’s t-test).

### Dot/Icm mediates lipid mixing between cells and artificial liposomes

Lipid mixing could depend on the T4SS interacting with target membrane lipids alone, or with lipids and an unknown receptor. To address this question, we assayed lipid mixing between *L. pneumophila* and artificial liposomes composed of lipids alone. Liposomes were labeled with a FRET pair of covalently attached fluorescent lipids: NBD-phosphatidylethanolamine (NBD-PE; donor) and lissamine rhodamine-PE (LR-PE; acceptor). When present at sufficiently high density in the same membrane, NBD fluorescence emission is quenched by LR-PE. Upon fusion with an unlabeled membrane, the dyes become diluted, disrupting energy transfer and resulting in increased NBD fluorescence emission (dequenching), a direct readout of lipid mixing^46^. We incubated labeled liposomes with wild-type or mutant strains lacking specific Dot/Icm components and monitored NBD fluorescence emission as a function of time (Fig. 6A). Consistent with the cell-based assay results, wild-type *L. pneumophila* induced an increase in NBD signal, indicating efficient lipid mixing. Notably, the ATPase-deficient DotB_E191K_ mutant exhibited similar activity, confirming that ATP hydrolysis is dispensable for lipid mixing. The Δ*dotA* strain, which assembles only the OMCC, also promoted lipid mixing, suggesting that the OMCC alone is sufficient to engage artificial target lipid membranes. By contrast, lipid mixing was not observed in strains lacking DotG (Δ*dotG*) and restored when DotG was ectopically expressed in a Δ*T4* background. Strains expressing only the cytosolic ATPase DotO, or that lacked the Dot/Icm system (Δ*T4*), failed to promote lipid mixing (Fig. 6B). These findings confirm that DotG is the minimal essential factor for T4SS-dependent lipid mixing. Further, they show that DotG can mediate lipid mixing with a target membrane composed of lipids alone.

**Figure 6.**
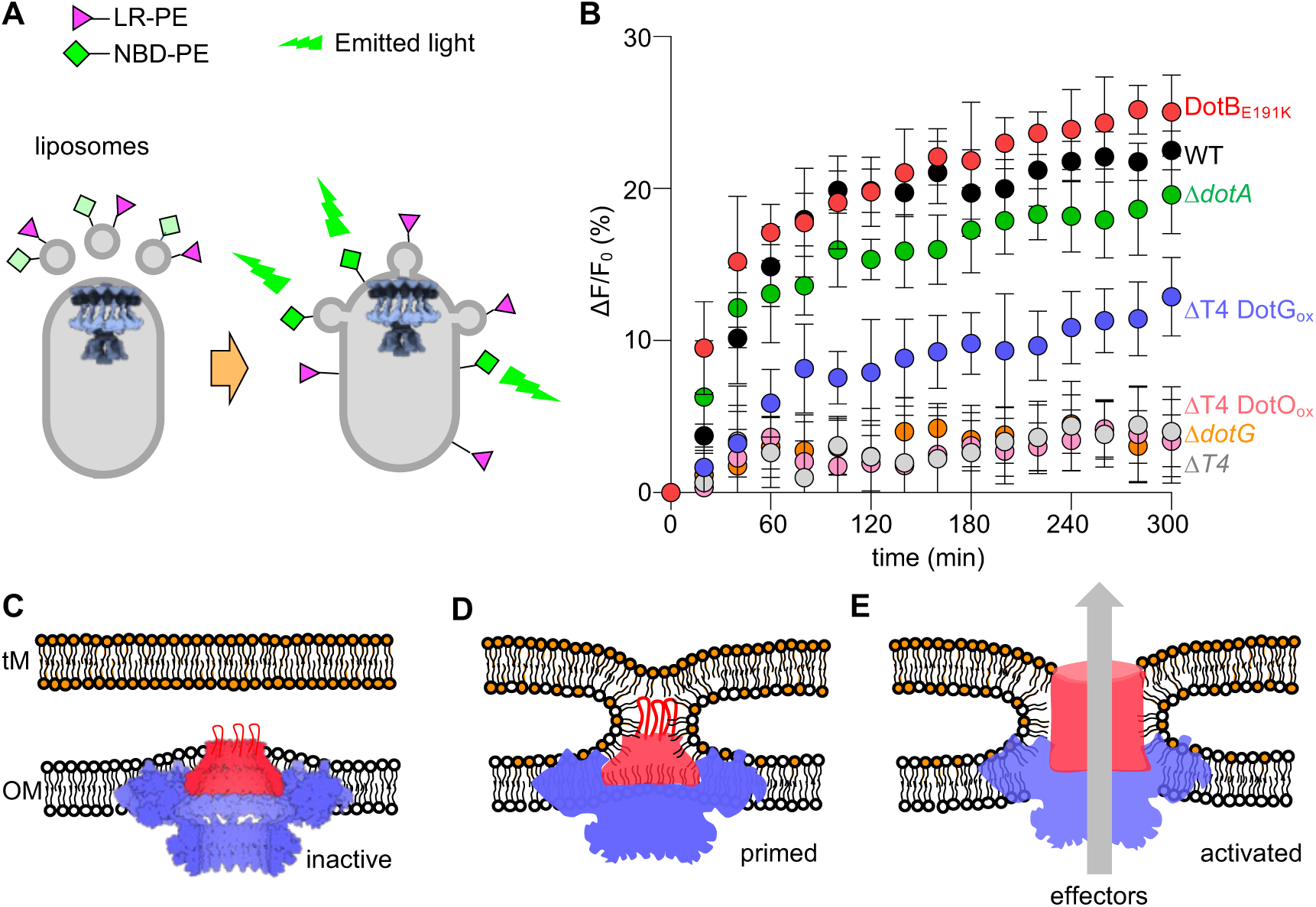
DotG-mediated lipid mixing with target membranes primes the Dot/Icm secretion system for activation. **(A)** Unlabeled *L. pneumophila* cells were incubated with liposomes containing a FRET lipid pair, NBD-phosphatidylethanolamine (NBD-PE, green, donor) and lissamine rhodamine-phosphatidylethanolamine (LR-PE, pink, acceptor). Lipid exchange between bacterial membranes and liposomes dilutes the FRET pair, reducing energy transfer and leading to dequenching of NBD-PE fluorescence. **(B)** NBD-PE fluorescence emission at 536 nm (excitation = 460 nm) was recorded for each condition, after mixing liposomes with bacteria. Strains tested were wild type (WT), a complete Dot/Icm deletion mutant (Δ*T4*), a DotA deletion mutant (Δ*dotA*), an ATPase-locked DotB_E191K_ mutant that assembles a complete but inactive apparatus, and a DotG deletion mutant (Δ*dotG*) that assembles only the outer membrane core complex (OMCC) lacking DotG. Complementation was achieved by plasmid expression of DotG or DotO in the ΔT4 background (ΔT4 DotG_ox_ and ΔT4DotO_ox_). **(C-E).** Model of Dot/Im priming. Prior to contact, unstructured extracellular loops (red) of DotG extend toward the target membrane (tM), likely mediating the initial interaction (C). OMCC is shown in blue. Only the OMCC portion of DotG is shown (red). DotG extends along the fully assembled Dot/Icm complex all the way into the cytoplasm (not shown), so, it is ideally placed to transmit signals from its initial contact with the tM. Contact destabilizes the target membrane and leads to a primed state that is permissive to lipid transfer between the tM and the OM (D). We drew only the proximal leaflets in continuity, since this is likely easier to achieve; the actual structures are unknown. The loops of DotG extend further into the tM, pulling on DotG and signaling tM engagement to the rest of the complex. DotG-tM engagement (priming) allows downstream activation steps^74–76^, including the involvement of the cytosolic components such as the ATPase DotB, DotO the coupling protein DotL and the chaperones IcmS-W, leading to the formation of a translocation pore that enables effector proteins to be delivered into the host cell (E). Secreted DotA and IcmX (not shown) may contribute to priming and/or activation by further engagement with tM lipids and/or structural rearrangements at the interface.

## DISCUSSION

A central unresolved question is how the large, multi-component T4SS machines are primed for activation and efficient delivery of substrates across donor and target membranes. On the one hand, this process must be sufficiently rapid and specific: in the case of *L. pneumophila*, effectors delivered by the Dot/Icm T4BSS must be translocated directly into the host cytosol to evade degradation in lysosomes and enable intracellular replication. Premature or unregulated effector release would compromise both efficiency and specificity. On the other hand, the remarkable host range of *L. pneumophila*, which include amoebae, human macrophages, and even other bacteria, argues against a specific target membrane receptor for priming Dot/Icm. Structural studies of Dot/Icm in its non-translocating state show that the secretion channel is closed at the extracellular face of the apparatus, suggesting that a target membrane-triggered conformational change is required to open the translocation channel and allow effector passage^3,10,32^. However, the nature of this trigger has remained elusive. Our results show that contact with target membrane lipids alone is sufficient to prime Dot/Icm for downstream activation and effector translocation, which can explain several previous observations.

First, priming of Dot/Icm by target membrane lipids would explain the broad tropism of *L. pneumophila,* because lipids are common components of membranes. Second, no conserved protein or glycan receptors have been identified on target membranes. If lipids prime Dot/Icm, there would be no need for such receptors. Third, consistent with the lack of a stable pilus that could mediate the initial contact with the target membrane to prime Dot/Icm, tight contacts between Dot/Icm and target membranes have recently been visualized^12,47^. Unlike the previously reported “closed” structures ^5,10,48^ in which Dot/Icm was not in contact with a target membrane, the recent tightly membrane apposed Dot/Icm structures- are in the “open”^47^, post-priming conformation, consistent with our results that target membrane lipids prime Dot/Icm for substrate transfer.

Although contact with the target membrane could in principle lead to Dot/Icm priming without lipid transfer between the two membranes, lipid mixing appears to be a functional and conserved part of the priming mechanism for the following reasons. First, lipid mixing is contact- and Dot/Icm-dependent, without need for ATP hydrolysis or even a fully assembled secretion system. These suggest that lipid mixing is an early step that precedes effector translocation that does not require energy consumption. Second, lipid mixing was not restricted to *L. pneumophila*; it was also observed in *E. coli* carrying the RK2 conjugation plasmid, indicating it is part of a priming mechanism conserved across at least some T4A and T4B secretion systems. Third, in the *E. coli*/RK2 system where we could simultaneously monitor lipid and substrate transfer, we found lipid mixing sometimes occurred in the absence of substrate transfer, but every substrate transfer event was associated with lipid mixing, implying lipid mixing precedes or accompanies substrate transfer. Fourth, lipids known to inhibit or promote membrane fusion inhibited or promoted substrate transfer, suggesting continuity of at least the proximal leaflets of the donor and acceptor membrane lipid bilayers. Overall, these observations show that lipid mixing is a prerequisite for substrate delivery.

Dot/Icm lacks a stable pilus so the lipid mixing and substrate transfer we observe must be a consequence of direct membrane-membrane contact. RK2, a T4ASS, can form a pilus, but the role of a pilus in substrate transfer is debated. While one study suggested substrate transfer occurs through the pilus^49^, others showed that in several T4ASSs, mutants that fail to form pili still support secretion, indicating that pilus formation is not essential for substrate delivery^50–59^. Interestingly, whether or not a pilus is involved in substrate transfer, pili may still play a role in lipid mixing, as pili themselves are lipid-protein composites^60–63^. Structural studies of F and F-like pili revealed that each pilin subunit is assembled with a phospholipid molecule, forming a stoichiometric protein-lipid complex^63^. Likewise, *A. tumefaciens* T-pili were shown to incorporate defined phospholipids, including phosphatidylglycerol and phosphatidylcholine, that line the pilus lumen and contribute to its electrostatic properties^60,61^. These findings indicate that pili may also assist lipid exchange with the target membrane in addition to their other functions. Nevertheless, our finding on the role of intrinsic positive- or negative-curvature lipids in RK2 transfer suggest lipid mixing results from the formation of a membrane stalk intermediate through direct membrane contact.

Although we do not yet know the molecular mechanisms leading to lipid mixing and T4SS priming, we identified DotG as both necessary and sufficient for lipid mixing. DotG is the outermost component of Dot/Icm. As such, it is best positioned to make initial contact with the target membrane. Based on structural studies and modeling, flexible extracellular loops from DotG extend from the OMCC toward the target membrane^5,10,26,47,48^. These loops, are conserved across DotG homologs^64^, remain unresolved in cryo-ET structures, as expected from their predicted flexibility. We propose that upon target membrane contact, DotG undergoes conformational changes that induce lipid mixing and simultaneously prime the secretion system for substrate transfer (Fig. 6C-E).

While DotG is required to initiate lipid mixing, subsequent steps must involve additional membrane rearrangements to enable effector passage through the acceptor membrane, a process that involves further components of the Dot/Icm machinery. DotA and IcmX have recently emerged as central players in establishing the translocation channel. DotA and IcmX are unusual and essential components of the Dot/Icm machinery, long known to be secreted into the extracellular milieu in a T4SS-dependent manner in both *L. pneumophila* and *Coxiella burnetii*^65–67^ (see also Fig S1). Although their functions were unclear, recent *in situ* structural studies revealed that DotA and IcmX assemble together as a pentameric protochannel at the central axis of the Dot/Icm machine, where they act as a gatekeeper for effector passage^47,68^. In the inactive state this channel is closed, preventing premature secretion, whereas upon activation it undergoes large-scale rearrangements to form an extended conduit across the bacterial envelope. Importantly, the assembly and secretion of DotA and IcmX require both a fully assembled Dot/Icm complex and ATPase activity, distinguishing their role from the earlier lipid mixing step we describe here.

Our results show that DotG-mediated target membrane engagement represents an initial priming event that occurs independently of ATPase activity. This engagement involves lipid exchange that is sensitive to positive- or negative-curvature lipids are known to modulate membrane fusion. Thus, it is possible that DotG-target membrane engagement leads to the merger of at least the proximal leaflets of the apposed membranes. An alternative route for lipid exchange would involve lipids moving between Dot/Icm subunits that must expand radially to allow large structural changes that are required for the formation of a central conduit in the activated structure. The formation of the central pore may involve release of DotA and IcmX that are initially plugging the center of the inactive Dot/Icm structure^69^. In either case, lipid mixing should occur before, and possibly during, DotA and IcmX release that remodels the secretion machine into an open state that is competent for effector translocation.

In summary, our data support a model for Dot/Icm priming and activation that is shown in Fig. 6C-E. First, contact between DotG and the target membrane destabilizes the target membrane and leads to a primed state that is permissive for lipid exchange, likely by establishing continuity between the proximal leaflets. Engagement of DotG with the target membrane is signaled to the rest of the complex. This signaling leads to the formation of a translocation pore that enables effector proteins to be delivered into the host cell. This likely involves release of DotA and IcmX. These components may contribute to priming and/or activation by further engagement with target membrane lipids and/or structural rearrangements at the interface.

## ACKNOWLEDGEMENTS

We thank all members of the Karatekin and Roy labs, in particular Ane Landajuela for valuable discussions and suggestions. This work was supported by the National Institutes of Health (NIH) grants R01NS122388 (to EK) R37AI041699 (to CR). The content is solely the responsibility of the authors and does not necessarily represent the official views of the NIH.

## AUTHOR CONTRIBUTIONS

All authors designed the research. DC performed all experiments, analyzed data with input from EK, and wrote the first draft of the manuscript with input from EK. All authors contributed to the final version.

## MATERIALS AND METHODS

### Bacterial strains and plasmids

Bacterial strains used in this study are derivatives of *L. pneumophila* strain (LP01) and *E. coli* Mach1, as listed in Table S1. *L. pneumophila* strains were cultured on charcoal yeast extract (CYE) agar plates at 37 °C, as previously described^70^. *E. coli* Mach1 strains were grown on LB agar plates supplemented with the appropriate antibiotics at 37 °C. For conjugation and lipid mixing assays, *L. pneumophila* donor cells harbored the non-conjugative plasmid pJB1806^71^, while acceptor cells expressed cytosolic GFP from the same plasmid to enable visualization. *E. coli* donor strains carried the conjugative RK2 plasmid (a T4ASS) encoding GFP at the AseI restriction site, permitting detection of substrate translocation. *E. coli* acceptor cells containing pJB1806 encoding GFP served as a negative control for plasmid transfer.

### *E. coli* conjugation

Overnight cultures of donor and recipient strains were grown in the presence of the appropriate antibiotics. Cultures (1 mL) were pelleted by centrifugation, washed once with PBS, and resuspended in the same buffer. Donor and recipient cells were then mixed at a 1:1 ratio (50 µL each), and 100 µL of the mixture was spotted onto a non-selective agar plate without spreading. For conjugation assays performed in the presence of oleic acid (OA) or lysophosphatidylcholine (LPC), the cell mixtures were incubated for 1 hour on non-selective plates prior to transfer to selective media. All conjugation plates were incubated overnight.

### Cell-cell lipid Mixing

Interbacterial lipid mixing assays with *L. pneumophila* were performed as follows: a 2-day heavy patch of donor cells was resuspended in water, and approximately 10⁵ donor cells were transferred into 1 mL of water. Donor cells were labeled with Di-8-ANEPPQ (Di8) or NileRed at a final concentration of 2.7 µM for 30 minutes at room temperature with occasional tube rotation to ensure uniform staining. Following incubation, cells were washed four times with water to remove excess dye, replacing the microcentrifuge tube after the first wash to minimize background fluorescence. Labeled donor cells were mixed with an equal number of unlabeled acceptor cells expressing cytosolic GFP. The mixture was centrifuged at 17,000 × g for 2 minutes, and the pellet was transferred onto a CYE agar plate. The plate was incubated at 37 °C for 2 hours to allow the cell pellet to adhere to the surface. Afterward, the cells were gently spread using an inoculating loop to facilitate expansion and incubated for an additional 16 hours at 37 °C prior to imaging. For *E. coli* assays, the same procedure was followed with the following modifications: donor and acceptor cells were collected from LB broth cultures, and prior to labeling, cells were washed with PBS to remove residual LB. NileRed was used as the fluorescent lipid dye at a final concentration of 10 µM, and all subsequent washing steps were performed using PBS. The final cell mixture was plated on LB agar.

### Cell-Liposome Lipid Mixing

Liposomes were prepared from a lipid mixture composed of 62% 1,2-dioleoyl-sn-glycero-3-phosphocholine (DOPC), 20% 1,2-dioleoyl-sn-glycero-3-phosphoethanolamine (DOPE), 15% 1,2-dioleoyl-sn-glycero-3-phospho-L-serine (DOPS), 1.5% NBD-phosphatidylethanolamine (NBD-PE), and 1.5% lissamine rhodamine-phosphatidylethanolamine (LR-PE). Lipids were mixed in chloroform, dried under a nitrogen stream, and further desiccated under vacuum for 2 hours. The dried lipid film was resuspended in water at a final total lipid concentration of 1 mM, vortexed thoroughly, and subjected to five freeze-thaw cycles. Unilamellar vesicles were then generated by extrusion through 0.1 µm polycarbonate membranes (Avanti Polar Lipids) using a mini-extruder. Two-day heavy patches of wild-type and mutant bacterial strains were suspended in water and washed. Bacterial cells were then incubated with the liposome suspension at a final concentration of 10^7^ cells/ml. Lipid mixing between bacterial membranes and liposomes was quantified using a Tecan Infinite M1000 fluorescence plate reader. NBD-PE fluorescence was measured at 20 minutes intervals for 360 minutes using λ_ex_=460 nm and λ*em*=536 nm. The increase in fluorescence was expressed as ΔF/F₀ over time.

### Fluorescent microscopy imaging and processing

Imaging of *L. pneumophila* and *E. coli* was carried out by resuspending 16 h patches in water or PBS, respectively. The cells were then spotted onto a thin pad of 1% agarose, covered with a coverslip, and immediately imaged at room temperature. Fluorescence micrographs were captured using a Nikon Eclipse TE2000-S inverted microscope equipped with a Spectra X light engine (Lumencor), a CoolSNAP EZ 20 MHz digital monochrome camera (Photometrics), and a Nikon Plan Apo 100× objective lens (1.4 numerical aperture), all controlled by SlideBook 6.0 software (Intelligent Imaging Innovations). Samples were imaged using 365 nm (120 mW), 485 nm (196 mW) or 560 nm (260 mW), LED excitation (Spectra X, Lumencor), with typical exposure times of 100–500 ms. Total fluorescence intensities were quantified using SlideBook by measuring the mean signal within bacterial cells, subtracting the background intensity, and multiplying the resulting value by the cell area. For *L. pneumophila* time-lapse micrographs (Fig. 2), the background region was defined using SlideBook. Image acquisition was carried out using minimal excitation intensities to minimize potential phototoxic effects. Fluorescence intensity measurements were corrected for non-specific photobleaching. The premise for the correction is that the overall intensity of non-bleached areas (of different cells in the same field) should remain constant over time. Intensity measurements were multiplied by the inverse of the ratio of fluorescence at a given time point over fluorescence at the initial time point.

### Immunofluorescence Microscopy

DotA immunofluorescence was done as previously described^72^. Briefly, one milliliter of *L. pneumophila* culture was added to 10 mL of 80% methanol and left for 1 hour at room temperature. The fixed cells were collected by centrifugation and resuspended in 1 mL of 80% methanol. The methanol-fixed cell suspension was dropped on a poly-L-lysine-coated slide and air-dried for 20 minutes. The dried sample was covered with lysozyme solution (2 mg/mL in 25 mM Tris-HCl [pH 8.0], 50 mM glucose, 10 mM EDTA) and incubated at room temperature for 10 minutes. The slide was then covered with PBST for 30 seconds and inclined to remove the solution. This step was repeated 3 times. The slide was covered with 100% methanol for 1 minute and inclined, then covered with 100% acetone for 1 minute and inclined. The slide was air-dried. PBST containing 2% bovine serum albumin (PBST-BSA) was placed on the slide to block the sample and removed after 15 minutes. The DotA monoclonal antibody mAb2.2948^73^ was applied to the slide and incubated for 1 hour at room temperature. The sample was then washed three times with PBST, followed by incubation with Alexa Fluor 555-conjugated anti-mouse secondary antibody together with Hoechst stain.

### Statistical analysis

All experiments were performed at least twice, unless otherwise noted. Differences in the distribution of total intensities were determined using a non-parametric test (Mann–Whitney), as most of samples displayed long tailed distributions. P < 0.05 was considered significant. For most microscopy experiments, roughly 1000 cells pers condition were analyzed. No statistical methods were used to predetermine sample size, and the researchers were not blinded to sample identity.

---

**Supplementary Table 1.** Strains used in this study.

| Name | Genotype | Description/comments | Source/reference |
| --- | --- | --- | --- |
| *Legionella pneumophila* strains |  |  |  |
| LPO1 | LPO1 | hsdR, Str<sup>R</sup> | 77 |
| $\Delta dotA$ | LPO1 $\Delta dotA$ | | 78 |
| wild type | LPO1 pJB1806 | hsdR, Str<sup>R</sup>, CM<sup>R</sup> | This study |
| $\Delta T4SS$ | LPO1 $\Delta T4SS$ pJB1806 | | This study |
| $\Delta dotA$ | LPO1 $\Delta dotA$ pJB1806 | | This study |
| $\Delta dotG$ | LPO1 $\Delta dotG$ pJB1806 | | This study |
| DotB<sub>E191K</sub> | LPO1 dotB<sub>E191K</sub> pJB1806 |  | This study |
| DotGox | LPO1 $\Delta T4SS$ pJB1806 | | This study |
| DotOox | LPO1 $\Delta T4SS$ pJB1806 | | This study |
| wild type-GFP | LPO1 pJB1806-GFP |  | This study |
| $\Delta T4SS$-GFP | LPO1 $\Delta T4SS$ pJB1806-GFP | | This study |
| $\Delta dotA$-GFP | LPO1 $\Delta dotA$ pJB1806-GFP | | This study |
| $\Delta dotG$-GFP | LPO1 $\Delta dotG$ pJB1806-GFP | | This study |
| DotB<sub>E191K</sub>-GFP | LPO1 dotB<sub>E191K</sub> pJB1806-GFP |  | This study |
| DotGox-GFP | LPO1 $\Delta T4SS$ pJB1806-GFP | | This study |
| DotOox-GFP | LPO1 $\Delta T4SS$ pJB1806-GFP | | This study |
| *Escherichia coli* strains |  |  |  |
| Mach1 | F- $\phi 80lacZ\Delta M15 \Delta lacX74$<br>hsdR(rK<sup>-</sup>, mK<sup>+</sup>) $\Delta recA1398$<br>endA1 tonA | Cloning strain | Thermo Fisher Scientific<br>Catalog #C862003 |
| RK2 | Mach1 pRK600 | CM<sup>R</sup> | This study |
| RK2-GFP | Mach1 pRK600 |  | This study |
| pJB1806 | Mach1 pJB1806 | CM<sup>R</sup> | This study |
| pJB1806-GFP | Mach1 pJB1806 |  | This study |
| phoP | K12 *phoP::kan* | Kan<sup>R</sup>, recipient in RK2<br>conjugation assay with<br>modulators | Kind gift from E.A.<br>Groisman<sup>79</sup> |

## REFERENCES

1. Isaac, D. T. & Isberg, R. Master manipulators: an update on Legionella pneumophila Icm/Dot translocated substrates and their host targets. Future Microbiol 9, 343–359 (2014).

2. Hubber, A. & Roy, C. R. Modulation of host cell function by Legionella pneumophila type IV effectors. Annu Rev Cell Dev Biol 26, 261–283 (2010).

3. Chetrit, D., Hu, B., Christie, P. J., Roy, C. R. & Liu, J. A unique cytoplasmic ATPase complex defines the Legionella pneumophila type IV secretion channel. Nat Microbiol 3, 678–686 (2018).

4. Segal, G. Identification of legionella effectors using bioinformatic approaches. Methods Mol Biol 954, 595–602 (2013).

5. Park, D., Chetrit, D., Hu, B., Roy, C. R. & Liu, J. Analysis of Dot/Icm Type IVB Secretion System Subassemblies by Cryoelectron Tomography Reveals Conformational Changes Induced by DotB Binding. mBio 11, e03328–19 (2020).

6. Kagan, J. C. & Roy, C. R. Legionella phagosomes intercept vesicular traffic from endoplasmic reticulum exit sites. Nat Cell Biol 4, 945–954 (2002).

7. Tilney, L. G., Harb, O. S., Connelly, P. S., Robinson, C. G. & Roy, C. R. How the parasitic bacterium Legionella pneumophila modifies its phagosome and transforms it into rough ER: implications for conversion of plasma membrane to the ER membrane. J Cell Sci 114, 4637–4650 (2001).

8. Vogel, J. P., Andrews, H. L., Wong, S. K. & Isberg, R. R. Conjugative transfer by the virulence system of Legionella pneumophila. Science 279, 873–876 (1998).

9. Luo, Z.-Q. & Isberg, R. R. Multiple substrates of the Legionella pneumophila Dot/Icm system identified by interbacterial protein transfer. Proc Natl Acad Sci U S A 101, 841–846 (2004).

10. Ghosal, D. et al. Molecular architecture, polar targeting and biogenesis of the Legionella Dot/Icm T4SS. Nat Microbiol 4, 1173–1182 (2019).

11. Nagai, H. et al. A C-terminal translocation signal required for Dot/Icm-dependent delivery of the Legionella RalF protein to host cells. Proc Natl Acad Sci U S A 102, 826–831 (2005).

12. Böck, D. et al. The Polar Legionella Icm/Dot T4SS Establishes Distinct Contact Sites with the Pathogen Vacuole Membrane. mBio 12, 10.1128/mbio.02180-21 (2021).

13. Costa, T. R. D. et al. Type IV secretion systems: Advances in structure, function, and activation. Mol Microbiol 115, 436–452 (2021).

14. Christie, P. J., Whitaker, N. & González-Rivera, C. Mechanism and structure of the bacterial type IV secretion systems. Biochim Biophys Acta 1843, 1578–1591 (2014).

15. Gonzalez-Rivera, C., Bhatty, M. & Christie, P. J. Mechanism and Function of Type IV Secretion During Infection of the Human Host. Microbiol Spectr 4, (2016).

16. Costa, T. R. D. et al. Secretion systems in Gram-negative bacteria: structural and mechanistic insights. Nat Rev Microbiol 13, 343–359 (2015).

17. Trokter, M., Felisberto-Rodrigues, C., Christie, P. J. & Waksman, G. Recent advances in the structural and molecular biology of type IV secretion systems. Curr Opin Struct Biol 27, 16–23 (2014).

18. Kubori, T. & Nagai, H. The Type IVB secretion system: an enigmatic chimera. Curr Opin Microbiol 29, 22–29 (2016).

19. Trokter, M., Felisberto-Rodrigues, C., Christie, P. J. & Waksman, G. Recent advances in the structural and molecular biology of type IV secretion systems. Curr Opin Struct Biol 27, 16–23 (2014).

20. Bhatty, M., Laverde Gomez, J. A. & Christie, P. J. The expanding bacterial type IV secretion lexicon. Research in Microbiology 164, 620–639 (2013).

21. Kolatka, K., Kubik, S., Rajewska, M. & Konieczny, I. Replication and partitioning of the broad-host-range plasmid RK2. Plasmid 64, 119–134 (2010).

22. Nagai, H. & Kubori, T. Type IVB Secretion Systems of Legionella and Other Gram-Negative Bacteria. Front Microbiol 2, 136 (2011).

23. Liu, X., Khara, P., Baker, M. L., Christie, P. J. & Hu, B. Structure of a type IV secretion system core complex encoded by multi-drug resistance F plasmids. Nat Commun 13, 379 (2022).

24. Khara, P., Song, L., Christie, P. J. & Hu, B. In Situ Visualization of the pKM101-Encoded Type IV Secretion System Reveals a Highly Symmetric ATPase Energy Center. mBio 12, 10.1128/mbio.02465-21 (2021).

25. Amin, H., Ilangovan, A. & Costa, T. R. D. Architecture of the outer-membrane core complex from a conjugative type IV secretion system. Nat Commun 12, 6834 (2021).

26. Sheedlo, M. J. et al. Cryo-EM reveals species-specific components within the Helicobacter pylori Cag type IV secretion system core complex. eLife 9, e59495 (2020).

27. Durie, C. L. et al. Structural analysis of the Legionella pneumophila Dot/Icm type IV secretion system core complex. eLife 9, e59530 (2020).

28. Hu, B. et al. In Situ Molecular Architecture of the Helicobacter pylori Cag Type IV Secretion System. mBio 10, 10.1128/mbio.00849-19 (2019).

29. Hu, B., Khara, P. & Christie, P. J. Structural bases for F plasmid conjugation and F pilus biogenesis in *Escherichia coli*. Proc. Natl. Acad. Sci. U.S.A. 116, 14222–14227 (2019).

30. Sgro, G. G. et al. Cryo-EM structure of the bacteria-killing type IV secretion system core complex from Xanthomonas citri. Nat Microbiol 3, 1429–1440 (2018).

31. Macé, K. et al. Cryo-EM structure of a type IV secretion system. Nature 607, 191–196 (2022).

32. Park, D., Chetrit, D., Hu, B., Roy, C. R. & Liu, J. Analysis of Dot/Icm Type IVB Secretion System Subassemblies by Cryoelectron Tomography Reveals Conformational Changes Induced by DotB Binding. mBio 11, e03328–19 (2020).

33. Li, Y. G. & Christie, P. J. The Agrobacterium VirB/VirD4 T4SS: Mechanism and Architecture Defined Through In Vivo Mutagenesis and Chimeric Systems. in Agrobacterium Biology: From Basic Science to Biotechnology (ed. Gelvin, S. B.) 233–260 (Springer International Publishing, Cham, 2018). doi:10.1007/82_2018_94.

34. Kirby, J. E., Vogel, J. P., Andrews, H. L. & Isberg, R. R. Evidence for pore-forming ability by Legionella pneumophila. Mol Microbiol 27, 323–336 (1998).

35. Tsau, Y. et al. Dye screening and signal-to-noise ratio for retrogradely transported voltage-sensitive dyes. J Neurosci Methods 70, 121–129 (1996).

36. Fluhler, E., Burnham, V. G. & Loew, L. M. Spectra, membrane binding, and potentiometric responses of new charge shift probes. Biochemistry 24, 5749–5755 (1985).

37. Clarke, R. J. & Kane, D. J. Optical detection of membrane dipole potential: avoidance of fluidity and dye-induced effects. Biochim Biophys Acta 1323, 223–239 (1997).

38. Skaliy, P. & McEachern, H. V. Survival of the Legionnaires’ disease bacterium in water. Ann Intern Med 90, 662–663 (1979).

39. Schofield, G. M. A note on the survival of Legionella pneumophila in stagnant tap water. J Appl Bacteriol 59, 333–335 (1985).

40. Hsu, S. C., Martin, R. & Wentworth, B. B. Isolation of Legionella species from drinking water. Appl Environ Microbiol 48, 830–832 (1984).

41. Waggoner, A. S. Dye indicators of membrane potential. Annu Rev Biophys Bioeng 8, 47–68 (1979).

42. Chernomordik, L. V. & Kozlov, M. M. Mechanics of membrane fusion. Nat Struct Mol Biol 15, 675–683 (2008).

43. Chernomordik, L. V. & Kozlov, M. M. Mechanics of membrane fusion. Nat Struct Mol Biol 15, 675–683 (2008).

44. Siegele, D. A. & Hu, J. C. Gene expression from plasmids containing the araBAD promoter at subsaturating inducer concentrations represents mixed populations. Proceedings of the National Academy of Sciences 94, 8168–8172 (1997).

45. Hui, C.-Y., Guo, Y., Zhang, W. & Huang, X.-Q. Rapid monitoring of the target protein expression with a fluorescent signal based on a dicistronic construct in Escherichia coli. AMB Express 8, 81 (2018).

46. Struck, D. K., Hoekstra, D. & Pagano, R. E. Use of resonance energy transfer to monitor membrane fusion. Biochemistry 20, 4093–4099 (1981).

47. Yue, J. et al. In situ structures of the Legionella Dot/Icm T4SS identify the DotA-IcmX complex as the gatekeeper for effector translocation. Proc Natl Acad Sci U S A 122, e2516300122 (2025).

48. Chetrit, D., Hu, B., Christie, P. J., Roy, C. R. & Liu, J. A unique cytoplasmic ATPase complex defines the Legionella pneumophila type IV secretion channel. Nat Microbiol 3, 678–686 (2018).

49. Goldlust, K. et al. The F pilus serves as a conduit for the DNA during conjugation between physically distant bacteria. Proceedings of the National Academy of Sciences 120, e2310842120 (2023).

50. Liu, X., Khara, P., Baker, M. L., Christie, P. J. & Hu, B. Structure of a type IV secretion system core complex encoded by multi-drug resistance F plasmids. Nat Commun 13, 379 (2022).

51. Banta, L. M. et al. An Agrobacterium VirB10 Mutation Conferring a Type IV Secretion System Gating Defect▿. J Bacteriol 193, 2566–2574 (2011).

52. Curtiss, R. Bacterial conjugation. Annu Rev Microbiol 23, 69–136 (1969).

53. Curtiss, R., Charamella, L. J., Stallions, D. R. & Mays, J. A. Parental functions during conjugation in Escherichia coli K-12. Bacteriol Rev 32, 320–348 (1968).

54. Haase, J., Lurz, R., Grahn, A. M., Bamford, D. H. & Lanka, E. Bacterial conjugation mediated by plasmid RP4: RSF1010 mobilization, donor-specific phage propagation, and pilus production require the same Tra2 core components of a proposed DNA transport complex. J Bacteriol 177, 4779–4791 (1995).

55. Jakubowski, S. J., Cascales, E., Krishnamoorthy, V. & Christie, P. J. Agrobacterium tumefaciens VirB9, an outer-membrane-associated component of a type IV secretion system, regulates substrate selection and T-pilus biogenesis. J Bacteriol 187, 3486–3495 (2005).

56. Jakubowski, S. J. et al. Agrobacterium VirB10 domain requirements for type IV secretion and T pilus biogenesis. Mol Microbiol 71, 779–794 (2009).

57. Jakubowski, S. J., Krishnamoorthy, V. & Christie, P. J. Agrobacterium tumefaciens VirB6 protein participates in formation of VirB7 and VirB9 complexes required for type IV secretion. J Bacteriol 185, 2867–2878 (2003).

58. Kishida, K. et al. Contributions of F-specific subunits to the F plasmid-encoded type IV secretion system and F pilus. Mol Microbiol 117, 1275–1290 (2022).

59. Sagulenko, E., Sagulenko, V., Chen, J. & Christie, P. J. Role of Agrobacterium VirB11 ATPase in T-pilus assembly and substrate selection. J Bacteriol 183, 5813–5825 (2001).

60. Amro, J. et al. Cryo-EM structure of the Agrobacterium tumefaciens T-pilus reveals the importance of positive charges in the lumen. Structure 31, 375–384.e4 (2023).

61. Kreida, S. et al. Cryo-EM structure of the Agrobacterium tumefaciens T4SS-associated T-pilus reveals stoichiometric protein-phospholipid assembly. Structure 31, 385–394.e4 (2023).

62. Vadakkepat, A. K. et al. Cryo-EM structure of the R388 plasmid conjugative pilus reveals a helical polymer characterized by an unusual pilin/phospholipid binary complex. Structure 32, 1335–1347.e5 (2024).

63. Costa, T. R. D. et al. Structure of the Bacterial Sex F Pilus Reveals an Assembly of a Stoichiometric Protein-Phospholipid Complex. Cell 166, 1436–1444.e10 (2016).

64. Tran, S. C., McClain, M. S. & Cover, T. L. Role of the CagY antenna projection in Helicobacter pylori Cag type IV secretion system activity. Infection and Immunity 91, e00150–23 (2023).

65. Nagai, H. & Roy, C. R. The DotA protein from Legionella pneumophila is secreted by a novel process that requires the Dot/Icm transporter. EMBO J 20, 5962–5970 (2001).

66. Luedtke, B. E., Mahapatra, S., Lutter, E. I. & Shaw, E. I. The Coxiella Burnetii type IVB secretion system (T4BSS) component DotA is released/secreted during infection of host cells and during in vitro growth in a T4BSS-dependent manner. Pathog Dis 75, ftx047 (2017).

67. Matthews, M. & Roy, C. R. Identification and subcellular localization of the Legionella pneumophila IcmX protein: a factor essential for establishment of a replicative organelle in eukaryotic host cells. Infect Immun 68, 3971–3982 (2000).

68. Yue, J. et al. In-situ structures of the Legionella Dot/Icm T4SS identify the DotA-IcmX complex as the gatekeeper for effector translocation. 2025.06.23.660953 Preprint at 10.1101/2025.06.23.660953 (2025).

69. Yue, J. et al. In situ structures of the Legionella Dot/Icm T4SS identify the DotA-IcmX complex as the gatekeeper for effector translocation. Proc Natl Acad Sci U S A 122, e2516300122 (2025).

70. Merriam, J. J., Mathur, R., Maxfield-Boumil, R. & Isberg, R. R. Analysis of the Legionella pneumophila fliI gene: intracellular growth of a defined mutant defective for flagellum biosynthesis. Infect Immun 65, 2497–2501 (1997).

71. Bardill, J. P., Miller, J. L. & Vogel, J. P. IcmS-dependent translocation of SdeA into macrophages by the Legionella pneumophila type IV secretion system. Molecular Microbiology 56, 90–103 (2005).

72. Hiraga, S., Ichinose, C., Niki, H. & Yamazoe, M. Cell Cycle–Dependent Duplication and Bidirectional Migration of SeqA-Associated DNA–Protein Complexes in E. coli. Molecular Cell 1, 381–387 (1998).

73. Matthews, M. & Roy, C. R. Identification and subcellular localization of the Legionella pneumophila IcmX protein: a factor essential for establishment of a replicative organelle in eukaryotic host cells. Infect Immun 68, 3971–3982 (2000).

74. Cascales, E. & Christie, P. J. Agrobacterium VirB10, an ATP energy sensor required for type IV secretion. Proc Natl Acad Sci U S A 101, 17228–17233 (2004).

75. Cascales, E., Atmakuri, K., Sarkar, M. K. & Christie, P. J. DNA substrate-induced activation of the Agrobacterium VirB/VirD4 type IV secretion system. J Bacteriol 195, 2691–2704 (2013).

76. Banta, L. M. et al. An Agrobacterium VirB10 mutation conferring a type IV secretion system gating defect. J Bacteriol 193, 2566–2574 (2011).

77. Berger, K. H. & Isberg, R. R. Two distinct defects in intracellular growth complemented by a single genetic locus in Legionella pneumophila. Mol Microbiol 7, 7–19 (1993).

78. Andrews, H. L., Vogel, J. P. & Isberg, R. R. Identification of linked Legionella pneumophila genes essential for intracellular growth and evasion of the endocytic pathway. Infect Immun 66, 950–958 (1998).

79. Zwir, I. et al. Dissecting the PhoP regulatory network of Escherichia coli and Salmonella enterica. Proc Natl Acad Sci U S A 102, 2862–2867 (2005).

