## Supplemental Figure 1 for "Type IV Secretion System Drives Lipid Mixing"

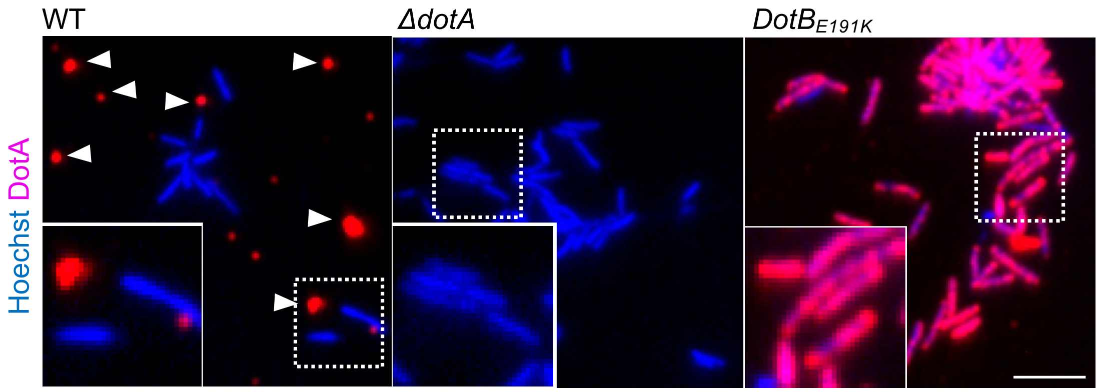


**Supplementary Figure 1. Secretion of DotA requires functional Dot/Icm ATPase activity.** Immunofluorescence micrographs showing the cellular localization of DotA (red) and DNA (Hoechst, blue) in WT, ATPase-deficient $DotB_{E191K}$, and ΔdotA mutants *L. pneumophila* strains. In WT cells, DotA is detected extracellularly (arrowheads), consistent with its secretion through a functional Dot/Icm system. DotA accumulates intracellularly in the $DotB_{E191K}$ mutant, indicating a block in secretion despite the presence of an assembled apparatus. Scale bar, 3 µm
